# Addressing the Elephant in the Room: A Quantitative Approach to Understanding “Wild”

**DOI:** 10.64898/2026.06.09.730786

**Authors:** Laura Saggiomo, Bruno Esattore, Janire Castellano Bueno, Michaela Masilkova, Andrea Sommese, Valeria Mazza, Sarah Gore, Pizza Ka Yee Chow

## Abstract

The concept of “wild” is poorly defined when applied to non-human animals, despite its central role in conservation, wildlife management and environmental policy. Despite numerous attempts to define “wildness”, the distinction between “wild” and “non-wild” remains inconsistently applied. A shared, empirically grounded understanding of wildness would strengthen communication, support policy and legislation, and clarify human attitudes towards wildlife and nature. We assessed whether a shared understanding of wildness exists within the research community. A total of 358 professionals from different fields completed an online survey comprising 44 wildness-related statements along established measures of attitude and acceptability toward wildlife (AATW) and nature-relatedness (NRS). . Exploratory Factor Analysis revealed six components (19 statements): Human-mediated animal availability, Urbanisation, Independence from humans, Human-wildlife perceived conflict, Individual history of wildness, and Feralisation. These factors explained 43.91% of the variance. Mediation analysis showed that urbanisation and human-wildlife perceived conflict shape nature-relatedness by influencing attitudes and acceptability towards wildlife. Respondents’ demographic and professional backgrounds influenced their conceptualisation of wildness. While there is a shared latent structure of wildness across fields, it is multidimensional. This has implications for developing working definitions applicable to research, policy and practice.

## Introduction

The concept of “wild” occupies a central place in several disciplines, influencing research, policy, and societal perceptions of nature. When applied to animals, the conceptual boundaries of “wild” carry substantial practical consequences for species protection and management. Whether a species is designated as “wild”, “semi-wild”, or “domestic” directly determines eligibility for conservation funding, IUCN Red List inclusion, and legal protection under wildlife legislation, with populations outside formal classifications at particular risk of exclusion from both management and welfare programmes [1–4]. However, the “wild” classification can also have negative implications when species are simultaneously categorised as pests or invasive, which can lead to subsequent management actions [5].

Despite numerous attempts to define what “wild” means (e.g., [6–8]), the distinction between “wild” and “non-wild” is still blurred and inconsistent. Human–nature relations, coupled with extensive land-use change, agriculture, urban development, wildlife management, and conservation practices, complicate efforts to draw clear boundaries between the two [9,10]. Defining “wild” also poses cultural challenges: conservation practice is largely shaped by Western knowledge systems, which often marginalise other perspectives, including Indigenous views [11]. Consequently, defining “wild” in a clear, fair, and consistent way has become increasingly complex. Beyond affecting policies, management, and conservation efforts, the conceptual boundaries of “wildness” carry a strong theoretical importance. For instance, this concept can affect people’s attitudes or reactions to the world [12] and, in particular, towards acceptance of wildlife [13–15] and relatedness to ‘nature’ [16].

Previous attempts at categorisations or discerning traits for animals to be labelled as “wild” or “not-wild” include definitions based on their physiological and behavioural traits, e.g., [17], morphological and molecular traits [18], along a wild-domesticated continuum [19] or, simply, a binary distinction between non-domesticated and domesticated [20]. Such inconsistency has limited comparability across fields, creating confusion that hinders interdisciplinary collaboration. The impacts of this confusion are particularly evident in fields related to animal behaviour, animal ecology, wildlife conservation, and animal welfare and management, where conceptual clarity has direct application relevance; a clear definition of “wild” is fundamental for deploying effective conservation actions [15,21], advancing environmental and social equity [22,23], shaping the ethical and cultural values assigned to species and populations [19], ensuring legal clarity [2], and facilitating inter-disciplinary communication and collaboration, to name a few.

Here, we used a quantitative, data-driven approach to examine the concept of “wild” across professional fields (e.g., Animal Behaviour, Animal Ecology, Conservation, Welfare, and Management). We first explored the latent factors of “wild” and then evaluated the extent to which demographic characteristics (e.g., employment sector, age, cultural affiliation) affect them. Finally, we examined how these factors were associated with participants’ attitudes towards, and the acceptability of, wildlife [13–15], as well as nature-relatedness (e.g., [16,24]).

## Materials and Methods

This study employed a cross-sectional survey design with a quantitative methodology. The project started in May 2025, and data were collected between August and December 2025. Participants were invited to complete an online survey on the Joint Information Systems Committee (JISC). Participants read the Participant Information Sheet, provided consent, completed demographic questions (Table S1), and rated the level of agreement with a list of 44 wildness statements (see S1) and 17 statements from two established psychometric scales (see below). All ratings were recorded on a Likert scale (see below). A total of 364 participants completed the survey, which took no more than 15 minutes. The full study adhered to the British Psychological Society Research Ethics guidelines. The study was approved by the Division of Psychology Ethics Committees, University of Chester (PC140825).

### Attitude and Acceptability Toward Wildlife Scale (AATW)

Metcalf and colleagues [15] developed AATW, which included 11 positive and negative statements: six statements measured attitude toward wildlife, whereas five statements measured acceptability toward wildlife. An example statement is “*Wildlife helps maintain the balance of nature*”, rated from 1 (‘*Completely disagree*’) to 7 (‘*Completely agree*’). Negative statements were reverse-scored before the total score was calculated. A higher score indicated a more positive attitude and acceptability. Cronbach’s α was 0.79 in this study.

### Natural Relatedness Scale (NR-6)

Nisbet and Zelenski [16] developed the NR-6 short psychometric scale to measure individual differences in connection to nature. Four statements assessed individuals’ identification with nature, their sense of connectedness with nature that may be expressed through spirituality, their awareness or subjective knowledge about the environment, and feelings of their connection with nature, and another two statements assessed self-identified needs in connection with nature and their awareness of nature and wildlife around environments. Two examples of the six statements are “*My relationship to nature is an important part of who I am*” and ‘*My ideal vacation spot would be a remote, wilderness area*’, rated from 1 (‘*Disagree strongly*’) to 5 (‘*Agree strongly*’). A higher score of total nature relatedness indicated an individual had a stronger connection to nature. Cronbach’s α was 0.67 in this study.

### Wildness statement

To understand the concept of wildness, we employed a nine-step approach [25–27]: 1) identify the domain/key elements of wildness and item generation; 2) assess face and content validity; 3) pre-test questions among the project team; 4) sampling and survey distribution; 5) item reduction; 6) factor extraction; 7) tests of dimensionality; 8) assess reliability; and 9) tests of validity.

The author team completed steps 1-3, during which members from different professional backgrounds generated statements independently. The statements were contested among the team members and subsequently revised to ensure face validity. The final set included statements broadly covering words and concepts related to wildness and wildlife, e.g., “*Animals stop being wild once they are tamed*” or ‘*Stray animals are wild*’, rated from 1 (‘*Completely disagree*’) to 7 (‘*Completely agree*’). Step 4 involved sampling and distributing the survey through professional networks, societies, mailing lists, scientific meetings, conferences, and events, and sharing it across social media platforms (see S5). We determined the sample size by using a response-to-item ratio between 5:1 and 10:1 to reduce item misclassification on the wrong factor [28].

### Data analysis

Data analysis was conducted using JAMOVI (Version 2.6). Six responses were excluded due to > 10% missing data across two or more scales [29], leaving 358 responses for subsequent analyses. The remaining missing data were imputed using scale means [30]. All inferential tests were two-tailed, with a significance level set at *p* ≤ 0.05. To explore the concept of wildness, we completed steps 5-9, which included item reduction, factor extraction, dimensionality testing, reliability assessment, and validity testing via Exploratory Factor Analysis (EFA).

Before conducting the EFA, correlations between the wildness statements (‘items’) were examined using Pearson’s correlation [31]; this aimed to ensure that items were moderately correlated but not highly correlated (i.e., 0.2 > r > 0.8). Highly correlated items suggest redundancy that could reduce discriminant validity and are typically removed to avoid multicollinearity and duplication [31]. No statements violated this criterion, so all items were retained for the EFA (see Table S2). Assumptions checked for the initial EFA indicated a factorable correlation matrix and adequate sample size (Bartlett’s test of sphericity: *χ*^2^_946_ = 3440.75, *p* < 0.001, KMO = 0.69) [32,33]. All 44 items were subjected to the initial EFA using Principal Axis factor extraction with an *oblimin* rotation to explore latent constructs through common variance. Factor loading was set at 0.4. Factor retention used several criteria beyond eigenvalue□> □1 and scree plot, including parallel analysis to improve accuracy and avoid over- or under-extraction (e.g., [28,34,35]). We removed statements that did not load on any factors, only had one statement loaded on a single factor, or a statement cross-loaded on two or more factors (removed a total of 23 statements); these statements indicated poor discriminant validity [28]. A factor with 3 or more statements has been suggested as the minimal acceptable level of reliability [28]; however, two-item factors were also retained if factor loading was > 0.6 and Spearman-Brown reliability was > 0.7, to avoid losing potentially meaningful constructs during early exploration.

We repeated these steps in a second EFA with 21 items, which was deemed the final analysis because 19 statements loaded onto a factor with no cross-loading issues. Factors were labelled based on the pattern found across items. Internal consistency was assessed using Cronbach’s α reliability test for factors with three or more items (factors 1-3) and Spearman-Brown reliability for two-item factors (factors 4-6). We also report the Root Mean Square Error of Approximation (RMSEA) fit index, with values < 0.05 indicating excellent fit [36], which provides a preliminary indication of the overall extracted factor structure. Two items with negative factor loadings were reverse-scored before the analysis (marked with “*” in Table 1); this aimed to align with the concept of that factor. The total scores for each factor were computed from participants’ responses to the corresponding items on the Likert scale. Table 3 shows the descriptive statistics for each factor.

**Table 1.**
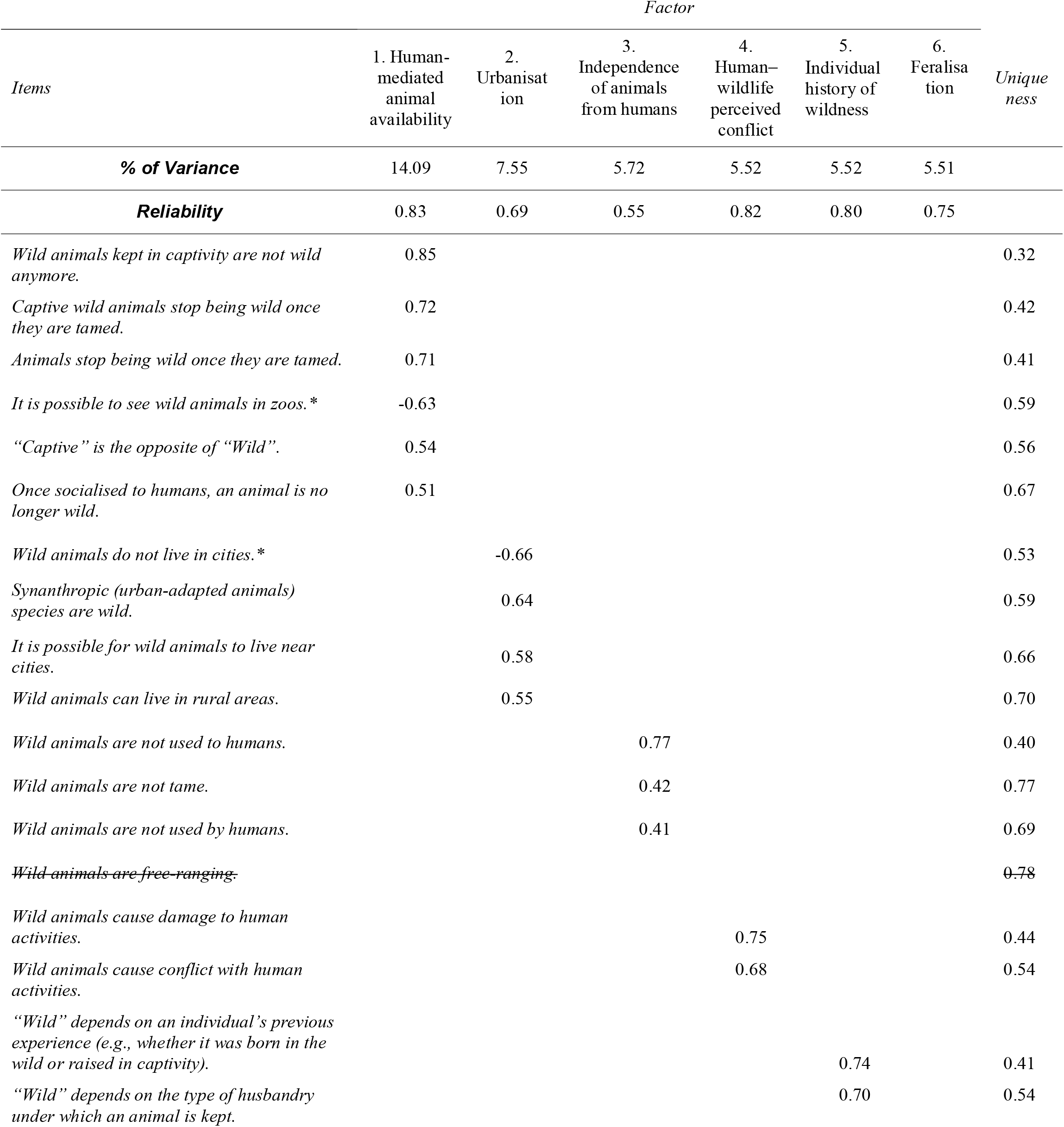
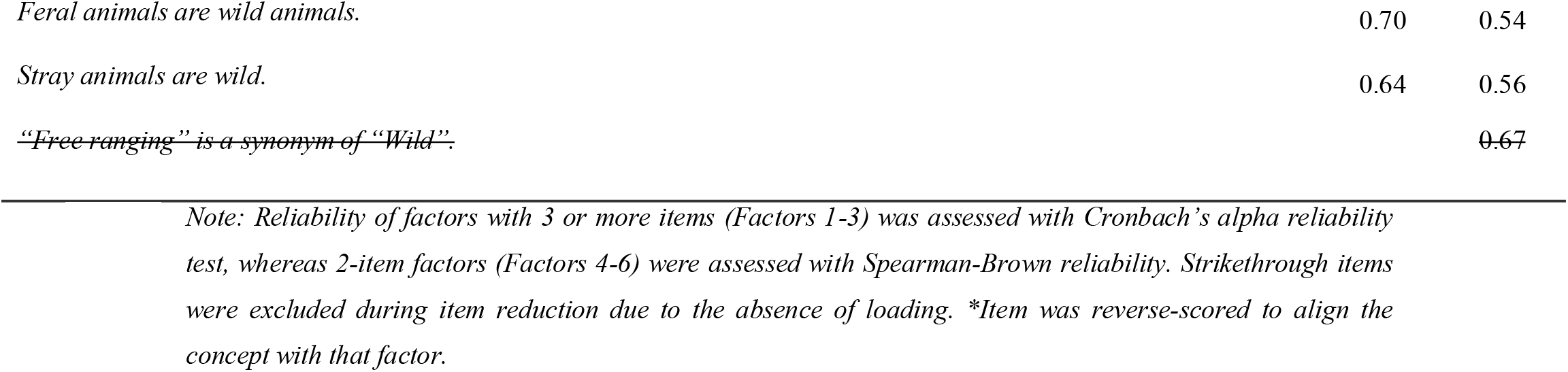
Exploratory factor analysis results: Factor loadings, uniqueness values, percentage of variance explained, and reliability coefficients from the final exploratory factor analysis (6-factor solution).

We ran a Generalised Linear Model (GLM) mediation analysis [37] to assess concurrent and predictive validity (Figure 1A), examining the direct effects of the six wildness factors on attitude/acceptance toward wildlife (AATW) and nature relatedness (NRS) and their indirect effects on nature relatedness through attitude/acceptance toward wildlife. Bootstrap resampling (5,000 iterations) provided 95% confidence intervals, given non-normal factor distributions (Shapiro-Wilks *p* < 0.05).

**Figure 1.**
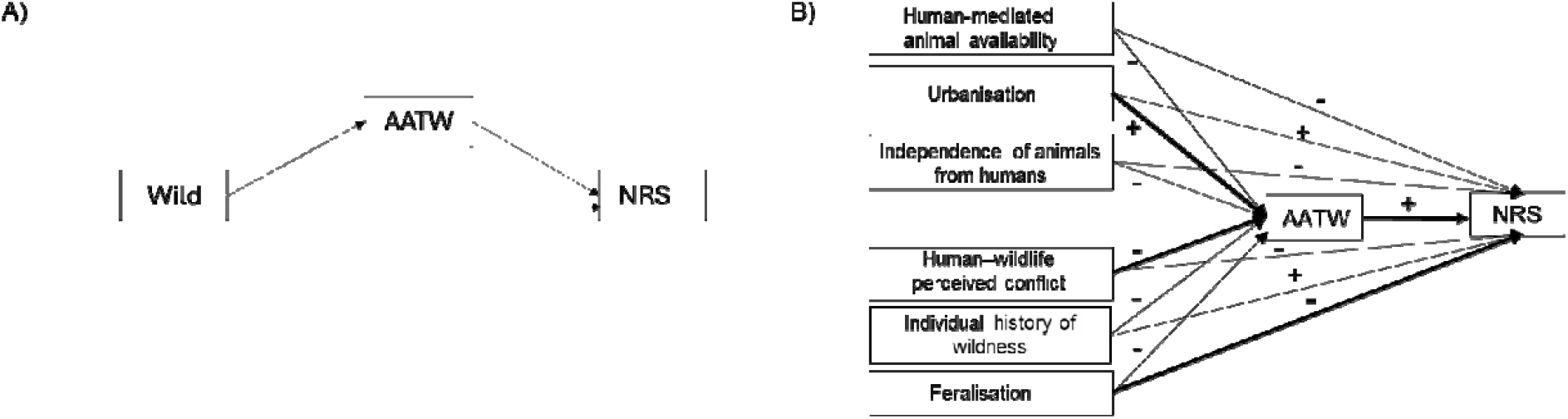
A) Conceptual model for assessing concurrent and predictive validity, proposing that wildness influences personal relatedness to nature (NRS) through its impact on attitude and acceptability toward wildlife (AATW). B) The mediation analysis result indicated the path direction and significance (p < 0.05 in bold) of the conceptual model.

Finally, we ran GLMs (‘GAMlj3’ package, [38]), examining the effects of demographic characteristics (Table S1) on each wildness factor. Predictors included age, sex, cultural affiliation, education level, number of years of animal-working experience, living area, and employment sector(s). All factors approximated normality except *Urbanisation*, which was log-transformed. All factors were modelled using a Gaussian identity link. The best model was selected via backward elimination *based on Akaike’s Information Criterion* (AIC) for each factor, with Pseudo-*R*^2^ and *χ*^2^ reported. Pairwise comparisons for significant categorical variables (> 2 levels) used the ‘emmeans’ package [39] with Tukey-adjusted *p*-values.

## Results

### Participants demographics

Full demographics are listed in Table S1. Participants (N = 358, *M* age = 38.78, *SD* = 11.96) (M = 42%, F = 58%) considered themselves working with wildlife or wild animals (mean working experience = 16.11 years, *SD* = 12.71), and engaged in multiple professional fields (*M* = 3.71, *SD* = 1.17), such as Animal Ecology, Animal Behaviour, and Animal Conservation. Fifty-seven percent of participants had a doctorate, 51% lived in an urban area, and the majority (94%) self-affiliated with Western culture.

### EFA: Item reduction, factor extraction, tests of dimensionality

As shown in Table 1, the final EFA included 21 statements to examine the underlying structure. This EFA met the assumption of sampling adequacy (KMO = 0.73) and sphericity (Bartlett’s test of sphericity: *χ*^2^_210_ = 1715.63, *p* < .001). Parallel analysis and scree plot inspection suggested retaining six factors (Figure S1), which together explained 43.91% of the total variance. There was no cross-loading issue, and factor loadings ranged from moderate to strong on each factor (0.41 - 0.85). These factors were weakly to moderately correlated (*r* = -0.13 to 0.25) (Table S2). Model fit indices indicated an acceptable solution (RMSEA = .05, *χ*^2^_99_ = 192.23, *p* < .001), reflecting good factor structure. Internal consistency of each factor was acceptable (Cronbach’s α = 0.55 – 0.83) (Table S3).

### Factor labelling and interpretation

The six factor labels were assigned based on the conceptual similarity shared among the items within that factor. Factor 1 ‘*Human-mediated animal availability*’ (6 items) reflected the role of humans in keeping animals available to them for their needs and wills. Factor 2 ‘*Urbanisation*’ (4 items) captured the significance of perceived spatial overlapping, or lack of it, between human infrastructures and wild animals’ habitats. Factor 3 ‘*Independence of animals from humans*’ (3 items) represented the perceived capability of animals to express their behavioural repertoire after contact with humans. Factor 4 ‘*Human–wildlife perceived conflict*’ (2 items) reflected the perceived negative side of wild animals’ presence on human activities. Factor 5 ‘*Individual history of wildness*’ (2 items) showed the fundamental role of human husbandry and breeding in determining an animal’s individual history of wildness. Finally, Factor 6 ‘*Feralisation*’ (2 items) captured the changes in human perception after wildlife acquired independence from humans.

### Concurrent and predictive validity

Full results of the mediation analysis are shown in Table S4. A higher attitude and acceptability toward wildlife (AATW) increased relatedness to nature (NRS) (Table 2, Figure 2B). Among the six wildness factors, a higher *Feralisation* score was directly associated with a lower NRS. The indirect effects of wildness factors on NRS were demonstrated through *Urbanisation*, and *Human–wildlife perceived conflict*. Urbanisation-related perceptions of animals had a significant positive effect on AATW, which in turn positively influenced NRS. This indicates that perceiving wildlife in urban environments is associated with more favourable attitudes towards wildlife, thereby strengthening individuals’ connection to nature. In contrast, *Human–wildlife perceived conflict* had a significant negative effect on AATW, which subsequently reduced NRS. Thus, higher levels of perceived conflict are associated with less favourable attitudes towards wildlife and a weakened sense of nature-relatedness.

**Table 2.**
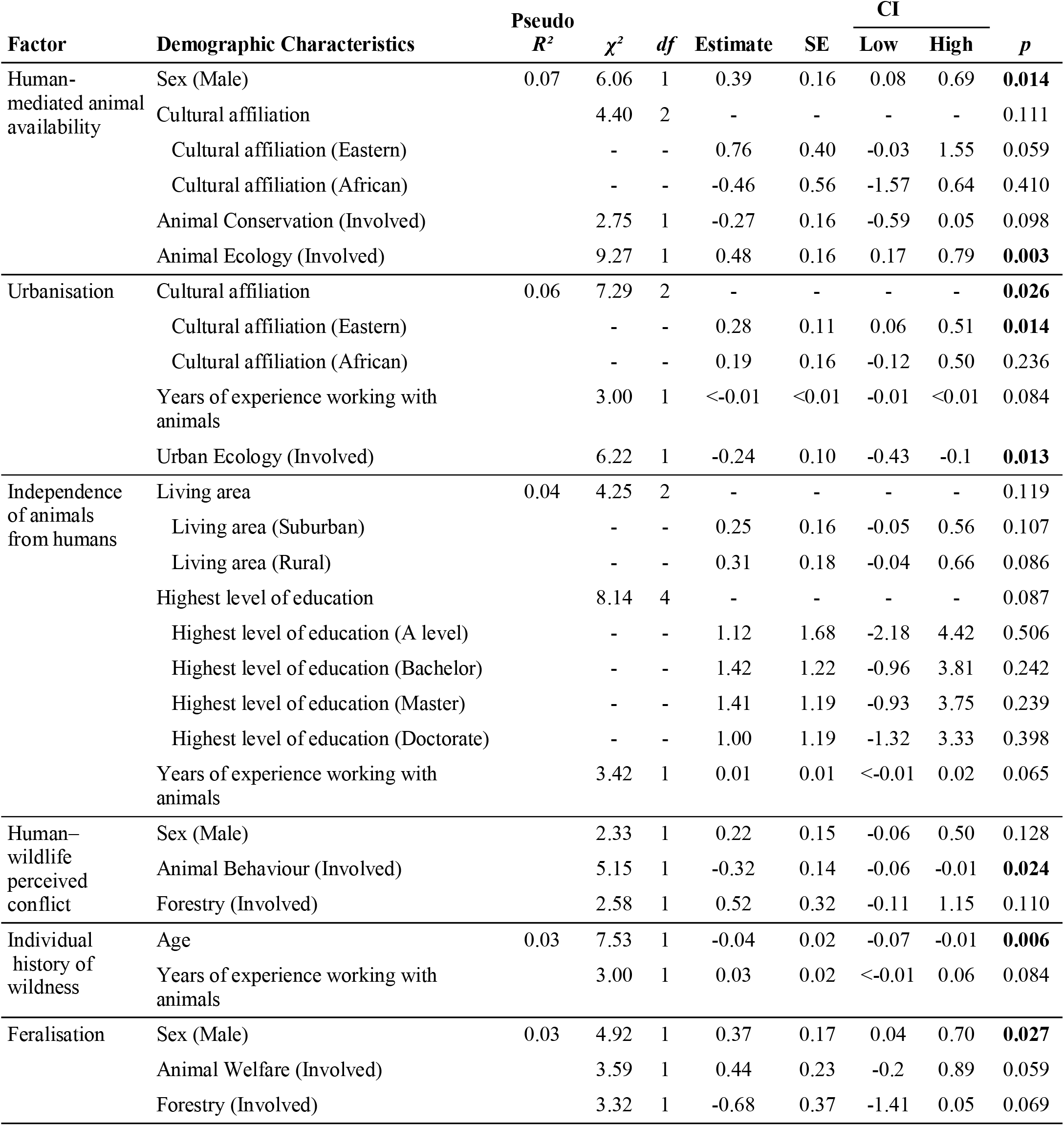
GLM results: parameter information of the ‘best’ models for each factor was selected using backward elimination based on Akaike’s Information Criterion (AIC). The reference group was ‘woman’ for sex, ‘western’ for cultural affiliation, ‘not-involved’ for employment sectors, ‘urban’ for living area, and ‘High school or equivalent’ for the highest level of education.

### Predictors for each of the extracted factors

Table 2 shows the best models selected via backward elimination using AIC explained 3-7% of the variance per factor. For *Human-mediated animal availability*, four predictors were retained (Table 2a), with sex and Animal Ecology being significant. Males were more likely than females to associate human mediation of animal accessibility with the concept of “wild”, and those working in the Animal Ecology sector were significantly more likely to do so compared with those not working in this sector. For *Urbanisation*, the best model included three predictors (Table 2b), with significant predictors being cultural affiliation and Urban Ecology. Eastern cultural affiliates, but not African affiliates, reported significantly higher agreement on this factor associated with “wild” than Western affiliates (Tukey-adjusted *p* = 0.038). Those who work inside Urban Ecology reported a lesser agreement than those outside. The best model for *Independence of animals from humans* retained three predictors (Table 2c), but none were significant predictors. For *Human–wildlife perceived conflict*, three predictors were retained (Table 2d), with Animal Behaviour being significant. Participants working in the Animal Behaviour sector are less likely to report that this factor is associated with the concept of ‘wild’. The best model of *Individual history of wildness* retained 2 predictors (Table 2e), with older participants being significantly less likely to agree that an individual’s history of being wild shapes the concept of ‘wild’. Finally, the best model of *Feralisation* included three predictors (Table 2f), with males being more likely than females to indicate that feralisation is associated with the concept of ‘wild’.

## Discussion

This project aimed to examine whether professionals across wildlife-related fields share a common underlying concept of “wild”, and if so, what structure that conceptualisation takes.

### Latent structure of “wildness”

Our results revealed six distinct factors of wildness: *Human-mediated animal availability, Urbanisation, Independence of animals from humans, Human-wildlife perceived conflict, Individual history of wildness*, and *Feralisation*. These factors, being only weakly to moderately intercorrelated, reflect that wildness is a multidimensional construct, in which each factor, despite varying degrees of relatedness, contributes a distinct understanding. The six factors explained approximately 44% of the variance, suggesting that this framework captures meaningful structure, with additional wildness-related dimensions that may exist and could be incorporated in future work to provide a more comprehensive account (e.g., welfare status, sentience).

These six factors can be organised around three broad conceptual themes: spatial confinement, degree of human intervention, and individual life experience, suggesting that researchers conceptualise wildness not as a binary opposition to domestication but as a multidimensional continuum. This structure aligns with the five-category framework proposed by Redford and colleagues [40], which distinguishes populations from self-sustaining wild to captive-bred along gradients of management intensity, suggesting that researchers’ implicit understanding of wildness aligns with existing theoretical classifications. The factors also echo philosophical distinctions related to dispositional wildness (tameness), constitutive wildness (being unadapted to human use), resource independence, and self-willed wildness e.g., [41,42], suggesting that these theoretically-derived categories have partial empirical support in researchers’ understanding of the concept. The factor structure also reflects a tension between individuals’ intrinsic value orientations, evident in factors concerned with animals’ independence and life experience, and more instrumental concerns, such as conflict and urban coexistence, though the structure leans overall toward intrinsic value dimensions [43].

### Validity of “wildness” factors

The mediation analysis demonstrated concurrent and predictive validity for the wildness construct; two of the wildness factors are associated with nature-relatedness (NRS), and this relationship is partially mediated by attitudes and acceptability toward wildlife (AATW), which is consistent with prior work linking wildlife attitudes to nature connectedness (e.g., [15,16]. More specifically, the wilderness factor *Urbanisation* increased AATW, and then NRS; this can be explained by respondents being able to recognise that wildlife can cohabit in urban spaces, and such recognition may be related to our innate affiliation with nature and wildlife that is stated in the Biophilia hypothesis [44]. The other wildness factor, *Human-wildlife perceived conflict*, negatively affected AATW and subsequently reduced NRS, reflecting that perceived risk and conflict increase negative attitudes toward wildlife and erode the human-nature connection [45–47]. Whilst not having a mediating role, we also found that *Feralisation* showed both a direct and an overall negative association with nature-relatedness, suggesting that perceiving animals as feral or de-wilded may independently diminish connectedness to nature, in parallel with or beyond attitude toward nature and wildlife, potentially because feral animals blur the categorical boundary between wild and domesticated.

### Demographic characteristics as predictors of “wildness” factors

Certain demographic characteristics stand out when examining the predictors of wildness factors, particularly respondents’ employment sectors or professional background. *Human-mediated animal availability* was predicted by Animal Ecology, suggesting that ecologists may hold a certain conception of what remains “wild” under human influence. *Urbanisation* was predicted by Urban Ecology, reflecting that discipline’s focus on how organisms adapt to urban habitats [48]. The weak positive intercorrelation between these two factors may reflect the conceptual overlap between Animal Ecology and Urban Ecology, as urbanisation can be understood as one form of human-mediated habitat modification. Finally, *Human-wildlife perceived conflict* was predicted by being involved in Animal Behaviour. This may partly reflect a disciplinary framing effect: research on human-wildlife interactions disproportionately focuses on conflict (71%), with minimal attention to coexistence (2%) or neutral interactions (8%) [49], potentially priming professionals in this field to associate wildness with its conflicting dimensions.

Other demographic characteristics that influence the conceptualisation of wildness factors include age, sex and cultural affiliation. *Individual history of wildness* was predicted by age, with younger respondents showing greater agreement that ‘wild’ is influenced by where an animal was born and the type of husbandry under which it is kept. This result can be related to younger respondents being more prone to have an individual-focused approach to wildlife than the older respondents, who tended to adopt a more management-oriented approach [50]. The fact that age was negatively associated with past wild animal experience may further reflect generational shifts in human-wildlife contact [50] due to human-driven habitat loss and fragmentation, which reduce the opportunity to encounter wildlife or have relatively less direct experience with wild animals in younger generations. *Human-mediated animal access* and *Feralisation* were predicted by sex. Sex differences in these two factors may reflect engagement in animal welfare, showing empathy, and human responsibility towards animals. It also reflects gender differences in wildlife-related experiences or risk perception [51,52], and could potentially be linked to documented differences in care-oriented attitudes toward individual animals [53]. Finally, *Urbanisation* was predicted by cultural affiliation, with Eastern cultural affiliates reporting higher agreement on perceived spatial overlap, or lack thereof, between human infrastructure and wild animals’ habitats than Western cultural affiliates. Although this finding should be interpreted cautiously due to the limited cultural diversity in the sample (i.e., 6%), it reveals a preliminary pattern in reflecting broader cultural differences and values in the perceived compatibility between wildlife and urban environments, potentially including a greater normalisation of urban-adapted animals as part of nature [54].

### Limitations and future direction

We made a first attempt to explore the underlying structure of wildness, and while the six factors explained approximately 44% of variance, results also suggest that the current items may not have captured the full construct of wildness. The unexplained variance likely reflects both individual experience and variation in how wildness is understood and conceptual dimensions not represented in the item list [55]. Future studies can incorporate a broader range of professional fields in item generation, which may help to explain variance not captured by the current six-factor solution and verify whether this structure is replicated across different disciplines. The observed intercorrelations among factors further suggest that the components of wildness are not fully independent, and the mediation analysis has only explored how wildness factors associate with some psychological and behavioural outcomes (i.e., AATW and NRS). Accordingly, the internal causal structure among the six factors themselves remains to be explored through theory-driven confirmatory work.

Additionally, our results showed that three of the six factors comprised only two items, which can limit their interpretive stability even though reliability was acceptable; two-item factors are sensitive to item-specific variance and require confirmation in independent samples. This highlights the next appropriate methodological step for this line of research is conducting confirmatory factor analysis (CFA), which allows assessing the construction and testing of theoretical models incorporating the psychological mechanisms linking wildness perception to conservation-relevant outcomes, such as attitude formation, perceived risk, and nature-relatedness, thereby strengthening the predictive utility of this framework for conservation practice and policy execution [14,56].

Our respondents are predominantly affiliated with a Western cultural background. Given that the perception of wildness does not exist in a static cultural or environmental context [54], our results likely have limited the generalisability of our findings to other cultures, and any observed cross-cultural differences should be treated as preliminary. “Wildlife” and “wild” do not translate uniformly across languages, and cultural-linguistic context likely shapes conceptualisation of wildness in ways this study could not fully capture [57]. The absence of equivalent terms in some languages may also reflect fundamentally different relationships between human communities and the natural world [58], which in turn likely affect ATTW and NRS, as well as how wildness is perceived and operationalised in research and policy e.g., [21,59,60]. A more culturally and linguistically diverse sample would strengthen both the generalisability of this framework and its utility for transboundary conservation efforts, where shared and translatable definitions are particularly important for international collaboration and trade agreements. Similarly, while the use of English enabled broad dissemination of the survey, it likely created a language barrier for many wildlife professionals with limited English proficiency, potentially reducing their likelihood of completing the questionnaire and contributing to sample bias.

Finally, ongoing anthropogenic habitat loss, fragmentation, and species range shifts mean that successive generations of researchers and practitioners are forming their understanding of wildness against an increasingly human-modified baseline, a process consistent with the concept of shifting environmental baselines in the Anthropocene (e.g., [61]). The negative association between age and *Individual history of wildness* observed here may partly reflect this dynamic, and longitudinal or cross-generational research would help clarify how perceptions of wildness are evolving [47].

## Conclusion

This study provides the first empirical, quantitative evidence that a shared concept of “wild” exists across professional fields, comprising six distinct but interrelated components: *Human-mediated animal availability, Urbanisation, Independence of animals from humans, Human-wildlife perceived conflict, Individual history of wildness, and Feralisation*. The weak to moderate intercorrelations among these factors confirm that the concept of “wild” is a multidimensional construct, organised around the degree and nature of human involvement in an animal’s life, environment, and history. Mediation analysis showed that the understanding of wildness can shape AATW and NRS. There are disciplinary patterns that suggest that the six-factor structure of wildness may be broadly latent across research communities but different fields selectively weight particular components more heavily and explicitly in practice, which may explain why a consensual, cross-disciplinary definition has remained elusive. Even within a discipline, experts’ personal characteristics may vary in their perceptions of wildness. Together, these findings offer a foundation for developing empirically grounded working definitions of “wild” that are not only of academic interest but also have implications, such as conservation communication and practice. While there is no single universal definition of wildness, researchers and practitioners may benefit from adopting a working definition that acknowledges wildness is multidimensional and explicitly specifies which components are in focus in a given project or policy context; this allows targeted conservation strategies to be communicated effectively, which can help minimise the perception of human-wildlife conflict and thereby human-wildlife conflicts themselves [62]. Future work requires confirmation of this structure through CFA, expanding cultural and linguistic representation, and tracking how wildness perceptions evolve across generations remain essential next steps.

## Acknowledgements

The authors would like to acknowledge all participants for their valuable contributions to this research.

