## Supplementary Materials for "Addressing the Elephant in the Room: A Quantitative Approach to Understanding “Wild”"

Preprint:

Affiliations:
[1]Swedish University of Agricultural Sciences; Department of Wildlife, Fish, and Environmental Studies, Sweden, 90183
[2] Wild Animal Initiative; 5123W 98th St, 1204, Minneapolis, MN 55437, USA
[3]Department of Game Management and Wildlife Biology, Faculty of Forestry and Wood Sciences, Czech University of Life Sciences Prague, Czech Republic, 16500
[4]Centre for Animal Nutrition and Welfare x Messerli Research Institute, University of Veterinary Medicine, Vienna (AT)
[5] Department of Ecological and Biological Sciences, University of Tuscia, Viterbo (VT), Italy, 01100
[6] Division of Psychology, University of Chester, Chester, UK CH1 3BJ

Content

Tables

S1 *Demographic characteristics of participants (N = 358).*
S2 *Inter-factor Pearson's r correlations.*
S3 *Summary of EFA results.*
S4 *Mediation analysis.*
S5 *Link to the list of professional contacts*

Figure

S1 *Parallel analysis on a scree plot of the final EFA result.*

**Table S1.** *Demographic characteristics of participants (N = 358). Note that the percentages are based on available data, as participants had the choice of not responding to demographic questions.*

| | N | % of Total |
|---|---|---|
| **Sex** | | |
| Female | 203 | 58 |
| Male | 146 | 42 |
| **Highest level of education** | | |
| High school or equivalent | 2 | 1 |
| A-levels degree | 3 | 1 |
| Bachelor's degree | 27 | 8 |
| Master's degree | 123 | 34 |
| Doctorate | 203 | 57 |
| **Living area** | | |
| Urban | 178 | 51 |
| Suburban | 99 | 29 |
| Rural | 70 | 20 |
| **Cultural affilitation** | | |
| Western | 316 | 94 |
| Eastern | 13 | 4 |
| African | 6 | 2 |
| **Employment sector(s) - multiple selections** | | |
| Animal Conservation | 133 | 17 |
| Animal Welfare | 57 | 7 |
| Animal Management | 65 | 9 |
| Animal Behaviour | 170 | 22 |
| Animal Ecology | 195 | 26 |
| Environmental Conservation | 76 | 10 |
| Forestry | 20 | 3 |
| Psychology | 18 | 2 |
| Urban Planning and Management | 4 | 1 |
| Urban Ecology | 23 | 3 |

**Table S2.** *Inter-factor Pearson's r correlations.*

| | 2. Urbanisation | 3. Independence of animals from humans | 4. Human–wildlife perceived conflict | 5. Individual history of wildness | 6. Feralisation |
|---|---|---|---|---|---|
| 1. Human-mediated animal availability | -0.13 | 0.25 | -0.05 | 0.24 | 0.13 |
| 2. Urbanisation | — | -0.16 | 0.01 | -0.12 | -0.00 |
| 3. Independence of animals from humans | | — | -0.05 | 0.03 | -0.02 |
| 4. Human–wildlife perceived conflict | | | — | 0.02 | -0.21 |
| 5. Individual history of wildness | | | | — | 0.14 |
| 6. Feralisation | | | | | — |

**Table S3.** *Summary of EFA results. Reliability of factors with 3 or more items (Factors 1-3) was assessed with Cronbach's alpha reliability test, whereas 2-item factors (Factors 4-6) were assessed with Spearman-Brown reliability.*

| **Factor** | ***SS Loadings*** | ***% of Variance*** | ***M(SD)*** | ***Reliability*** |
|---|---|---|---|---|
| 1. Human-mediated animal availability (6 items) | 2.96 | 14.09 | 3.41 (1.38) | 0.83 |
| 2. Urbanisation (4 items) | 1.58 | 7.55 | 6.37 (0.79) | 0.69 |
| 3. Independence of animals from humans (3 items) | 1.20 | 5.72 | 4.04 (1.18) | 0.55 |
| 4. Human–wildlife perceived conflict (2 items) | 1.16 | 5.52 | 4.40 (1.30) | 0.82 |
| 5. Individual history of wildness (2 items) | 1.16 | 5.52 | 3.46 (1.65) | 0.80 |
| 6. Feralisation (2 items) | 1.16 | 5.51 | 3.44 (1.45) | 0.75 |

**Table S4.** *Mediation analysis. Direct (D), Indirect (IN), Component (C) and total (T) effects (E) of wild concept on attitude and acceptability of wildlife and NRS. Confidence intervals computed with 5000 bias-corrected bootstrap. Estimates are standardised. Bold values are significant paths.*

| E | Path | | Estimate | SE | 95% C.I. (a) Lower | Upper | β | *z* | *p* |
|---|---|---|---|---|---|---|---|---|---|
| D | Human-mediated animal availability | ⇒ NRS | -0.09 | 0.11 | -0.30 | 0.13 | -0.04 | -0.79 | .429 |
| | Urbanisation | | 0.07 | 0.19 | -0.29 | 0.44 | 0.02 | 0.40 | .690 |
| | Independence of animals from humans | | -0.10 | 0.13 | -0.35 | 0.15 | -0.04 | -0.78 | .436 |
| | Human–wildlife perceived conflict | | -0.06 | 0.12 | -0.28 | 0.17 | -0.03 | -0.50 | .614 |
| | Individual history of wildness | | 0.08 | 0.09 | -0.10 | 0.25 | 0.05 | 0.87 | .382 |
| | Feralisation | | **-0.20** | **0.10** | **-0.39** | **-0.01** | **-0.11** | **-2.06** | **.040** |
| IN | Human-mediated animal availability | ⇒AATW ⇒NRS | -0.01 | 0.02 | -0.06 | 0.03 | -0.01 | -0.62 | .535 |
| | **Urbanisation** | | **0.14** | **0.05** | **0.04** | **0.25** | **0.04** | **2.69** | **.007** |
| | Independence of animals from humans | | -0.02 | 0.03 | -0.08 | 0.04 | -0.01 | -0.72 | .473 |
| | **Human–wildlife perceived conflict** | | **-0.13** | **0.04** | **-0.21** | **-0.05** | **-0.06** | **-3.28** | **.001** |
| | Individual history of wildness | | -0.00 | 0.02 | -0.04 | 0.03 | -0.00 | -0.16 | .872 |
| | Feralisation | | -0.01 | 0.02 | -0.05 | 0.04 | -0.00 | -0.31 | .757 |
| C | **AATW** | ⇒ **NRS** | **0.07** | **0.02** | **0.04** | **0.10** | **0.22** | **4.14** | **<.001** |
| | Human-mediated animal availability | ⇒AATW | -0.22 | 0.35 | -0.90 | 0.46 | -0.03 | -0.63 | .531 |
| | **Urbanisation** | | **2.07** | **0.58** | **0.93** | **3.22** | **0.18** | **3.55** | **<.001** |
| | Independence of animals from humans | | -0.30 | 0.41 | -1.09 | 0.50 | -0.04 | -0.73 | .466 |
| | **Human–wildlife perceived conflict** | | **-1.90** | **0.35** | **-2.60** | **-1.21** | **-0.27** | **-5.38** | **<.001** |
| | Individual history of wildness | | -0.05 | 0.28 | -0.60 | 0.51 | -0.01 | -0.16 | .872 |
| | Feralisation | | -0.10 | 0.31 | -0.71 | 0.52 | -0.02 | -0.31 | .756 |
| T | Human-mediated animal availability | ⇒ NRS | -0.10 | 0.11 | -0.32 | 0.12 | -0.05 | -0.91 | .365 |
| | Urbanisation | | 0.22 | 0.19 | -0.15 | 0.58 | 0.06 | 1.15 | .248 |
| | Independence of animals from humans | | -0.12 | 0.13 | -0.37 | 0.14 | -0.05 | -0.92 | .359 |
| | Human–wildlife perceived conflict | | -0.19 | 0.11 | -0.41 | 0.03 | -0.09 | -1.66 | .097 |
| | Individual history of wildness | | 0.07 | 0.09 | -0.10 | 0.25 | 0.04 | 0.82 | .413 |
| | **Feralisation** | | **-0.21** | **0.10** | **-0.41** | **-0.01** | **-0.11** | **-2.07** | **.038** |

**Table S5.** *List of professional contacts can be found in this link. This list includes contacts through professional networks, societies, mailing lists, scientific meetings, conferences, and events.*

**Contacts**

**Figure S1.** *Parallel analysis on a scree plot of the final EFA result.*

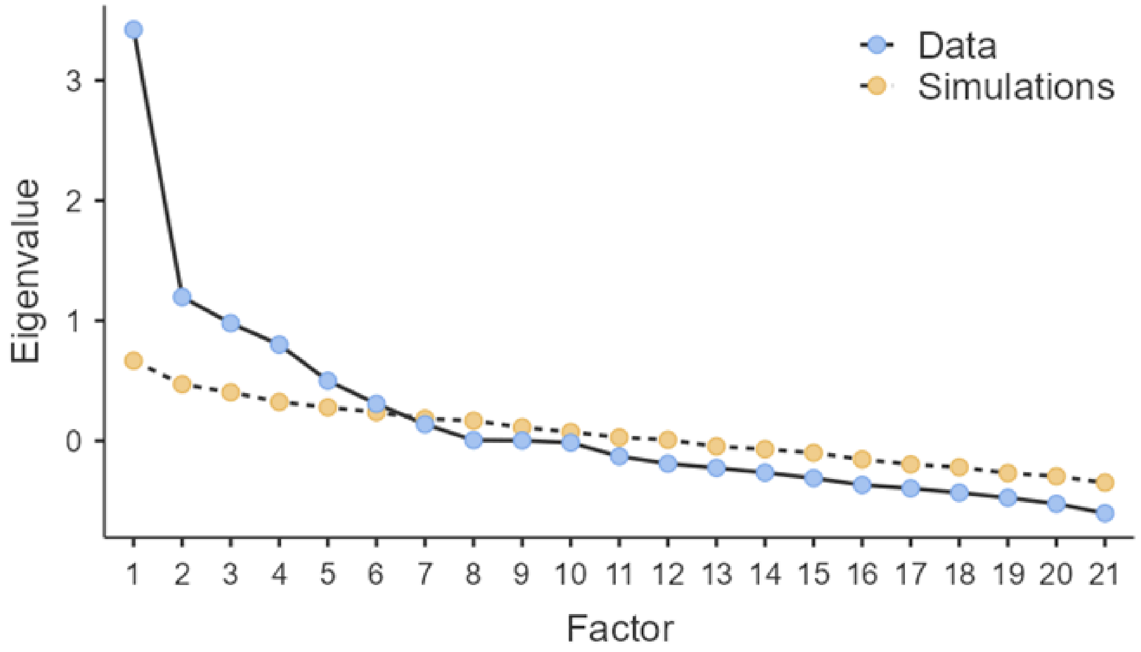
